## Supplementary figures and images for "The function of ER-phagy receptors is regulated through phosphorylation-dependent ubiquitination pathways"

### Supl Fig1

#### Supplementary figure 1

A

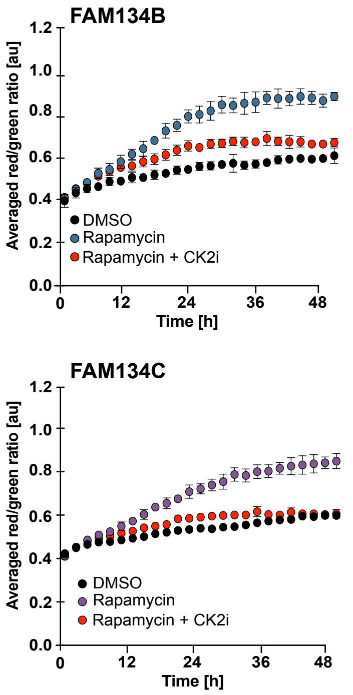


B

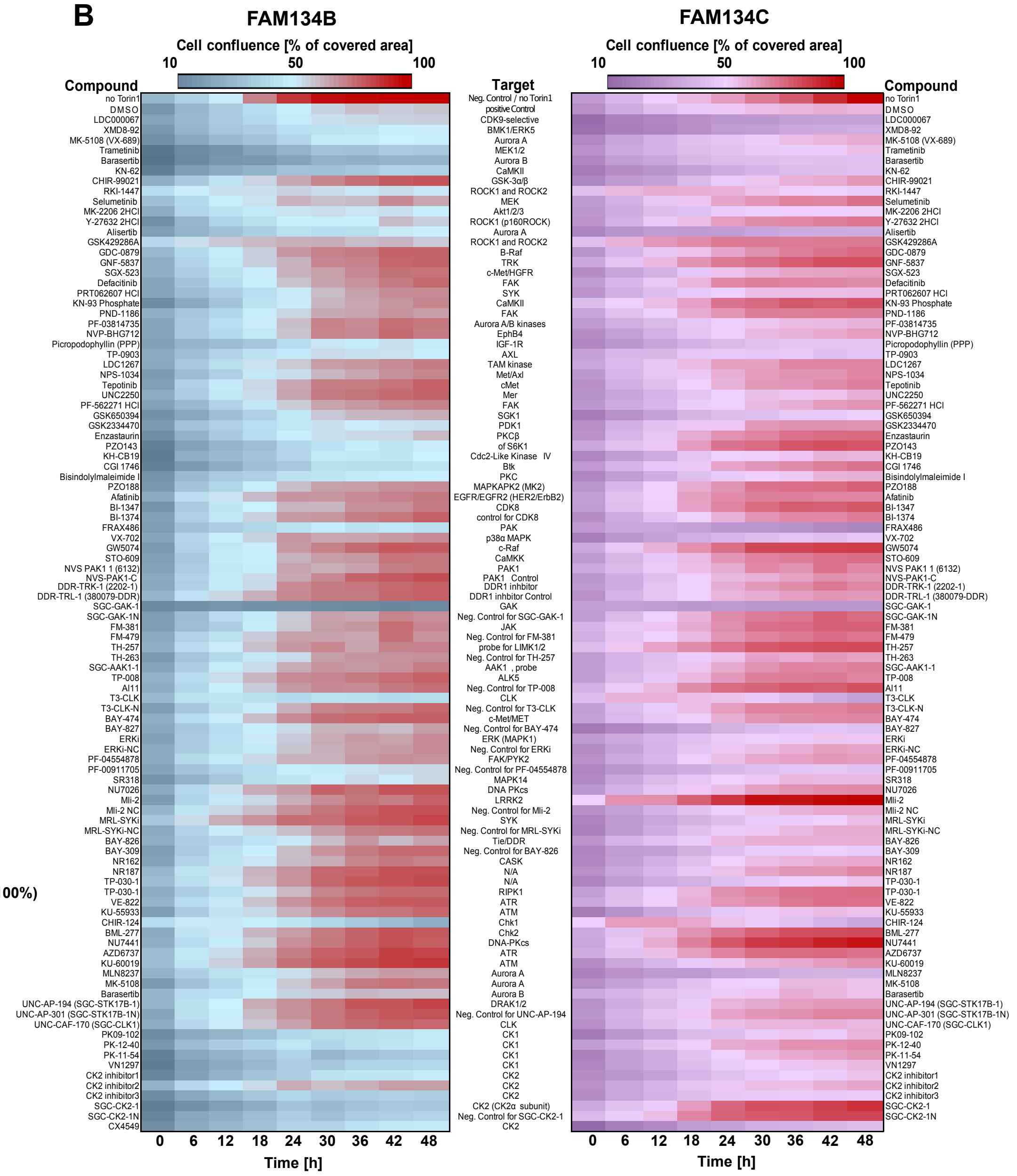


C

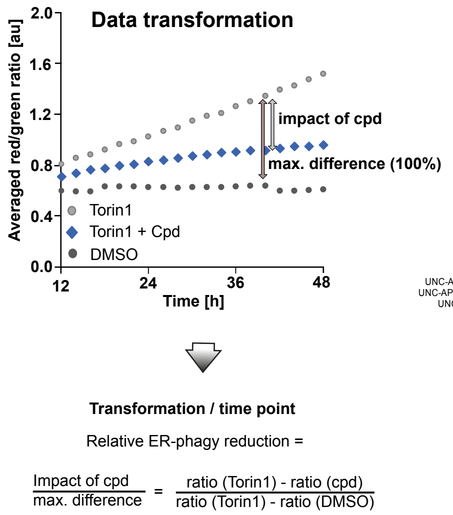


D

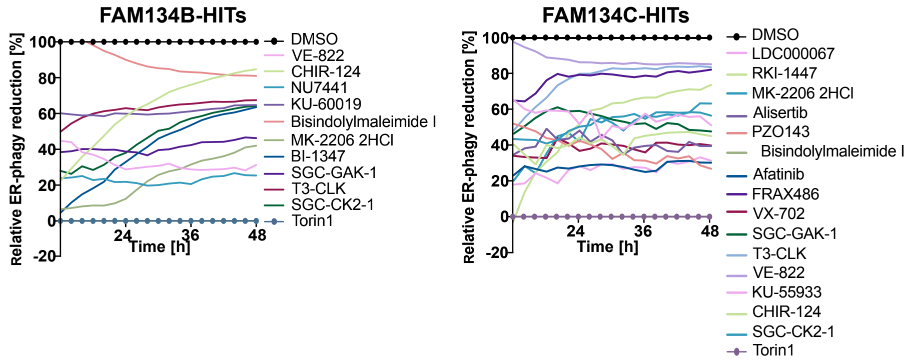

### Supl Fig4

#### Supplementary Figure 4

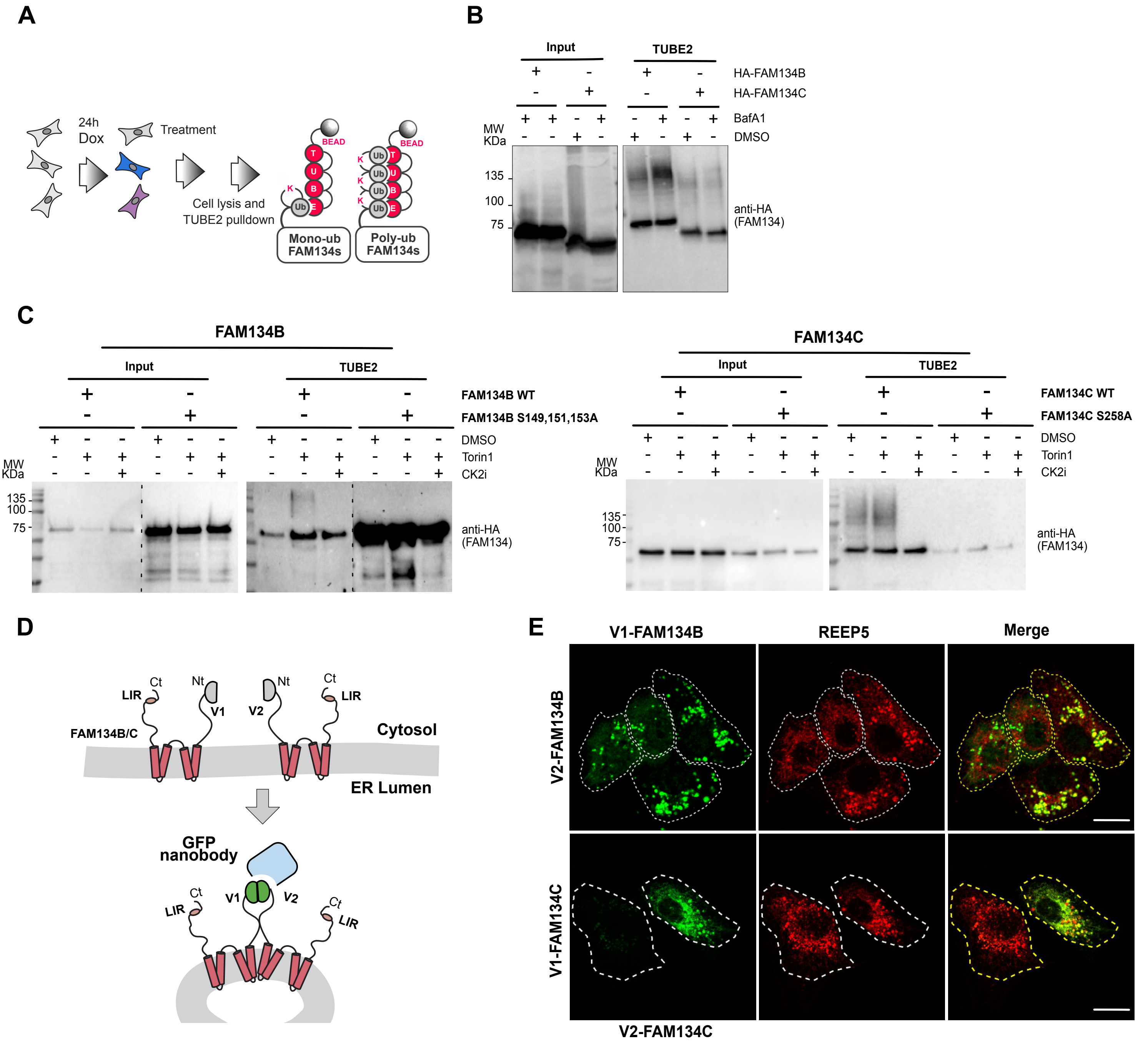

### Supl Fig5

#### Supplementary Figure 5

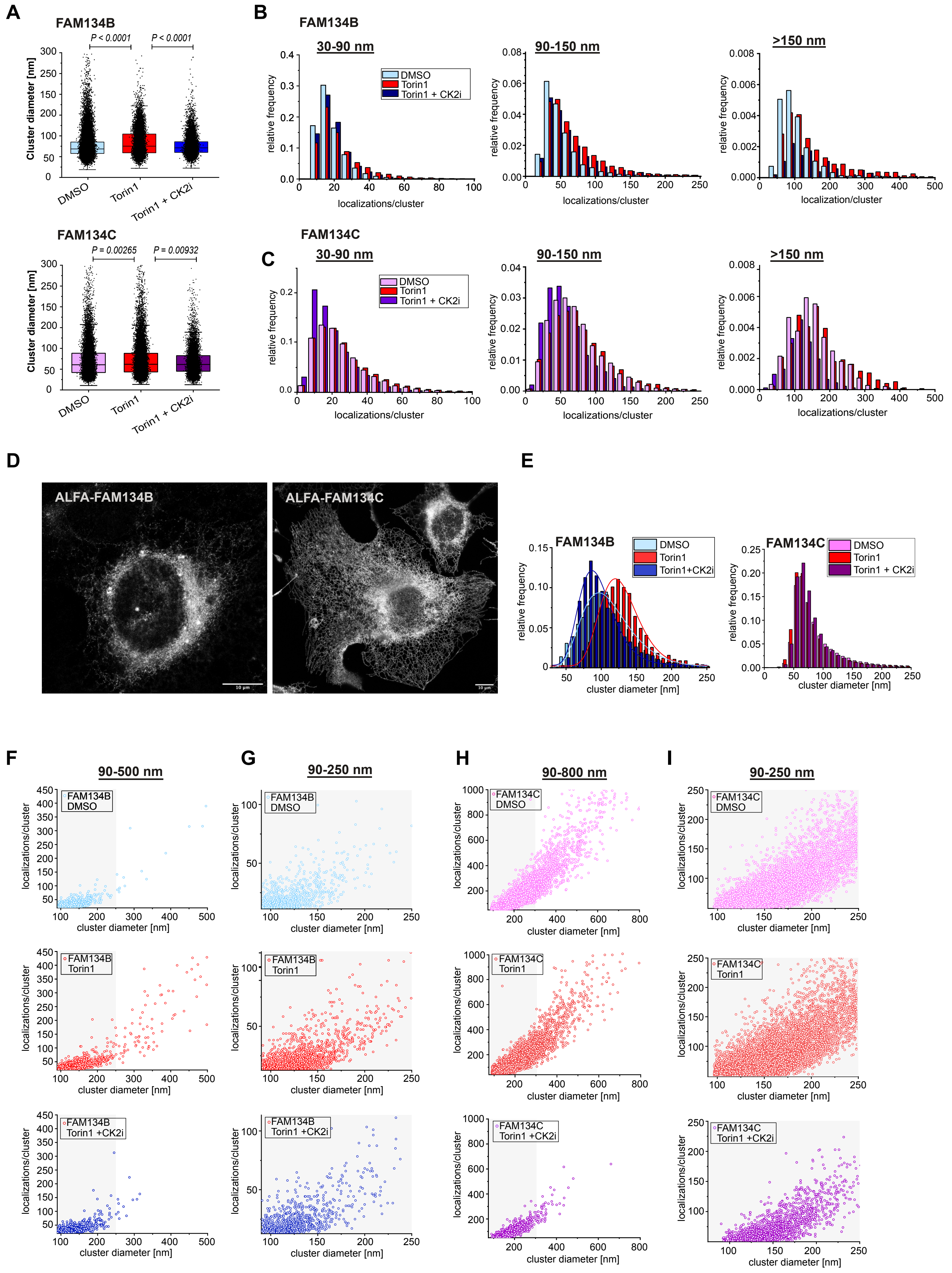

### Supl Fig6

#### Supplementary Figure 6

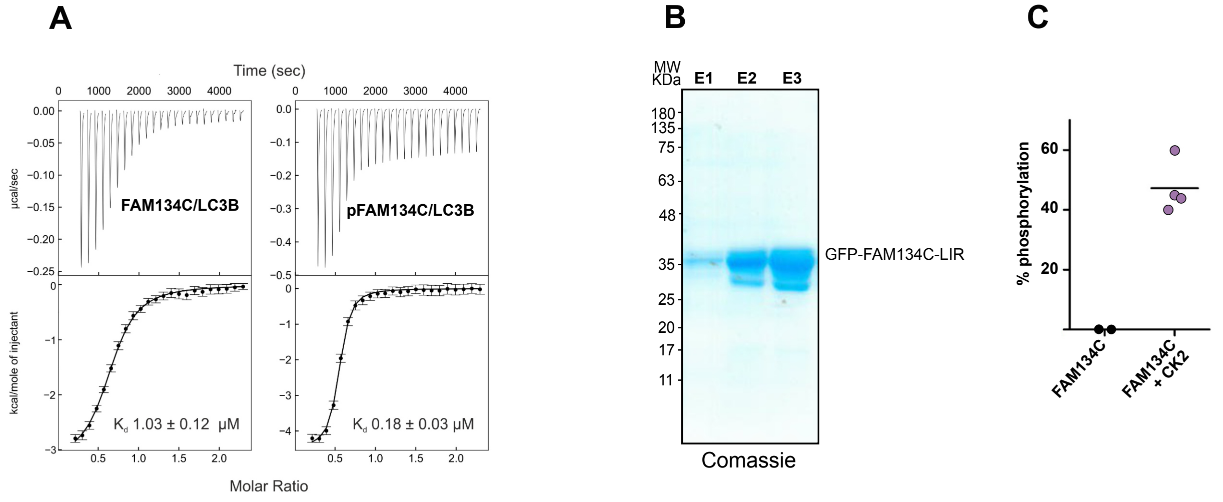


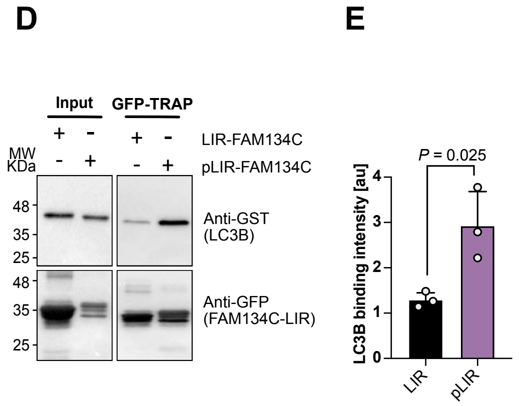
