## Supplementary material for "The function of ER-phagy receptors is regulated through phosphorylation-dependent ubiquitination pathways": Supl Fig2

### Supplementary Figure 2

**A**

**FAM134B-HIT list**

| Selected Hits with >20% Torin-1 induced ERphagy inhibition | Targeted kinase |
|---|---|
| SGC-CK2-1 | Catalytic subunits of CK2: CK2α (CSNK2A1) and CK2α' (CSNK2A2) |
| VE-822 | ATR |
| KU-55933 | ATM |
| CHIR-124 | Chk1 |
| MK-2206 2HCl | Akt1/2/3 |
| SGC-GAK1 | GAK |
| T3-CLK I | CLK |
| Bisindolylmaleimide I | PKC inhibitor I |
| NU7441 | DNA-PKCs |
| CHIR-99021 | GSK-3α/β |

**FAM134C-HIT list**

| Selected Hits with >20% Torin-1 induced ERphagy inhibition | Targeted kinase |
|---|---|
| SGC-CK2-1 | Catalytic subunits of CK2: CK2α (CSNK2A1) and CK2α' (CSNK2A2) |
| VE-822 | ATR |
| KU-55933 | ATM |
| CHIR-124 | Chk1 |
| MK-2206 2HCl | Akt1/2/3 |
| SGC-GAK1 | GAK |
| T3-CLK I | CLK |
| Bisindolylmaleimide I | PKC inhibitor I |
| Alisertib | Aurora A |
| VX-702 | p38α MAPK |
| FRAX486 | PAK |
| LDC0000067 | CDK9 |
| RKI-1447 | ROCK1 and ROCK2 |
| PZO143 | S6K1 |
| Afatinib | EGFR/EGFR2 (HER2/ErbB2) |

**B**

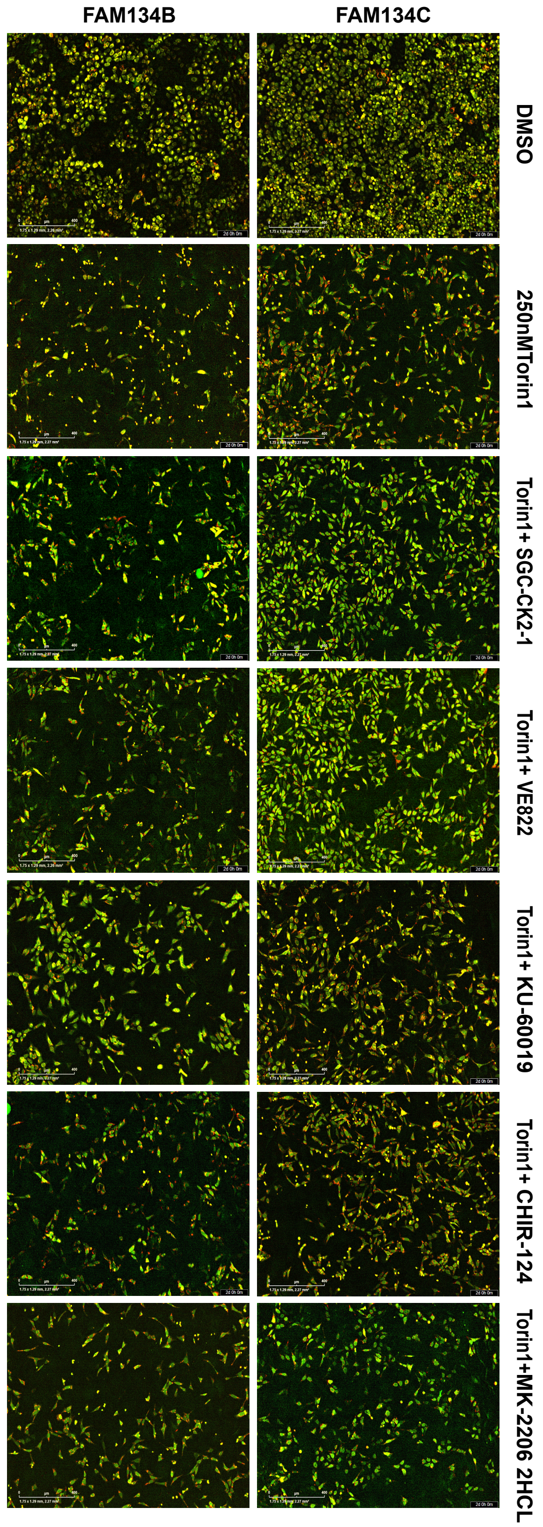


**C**

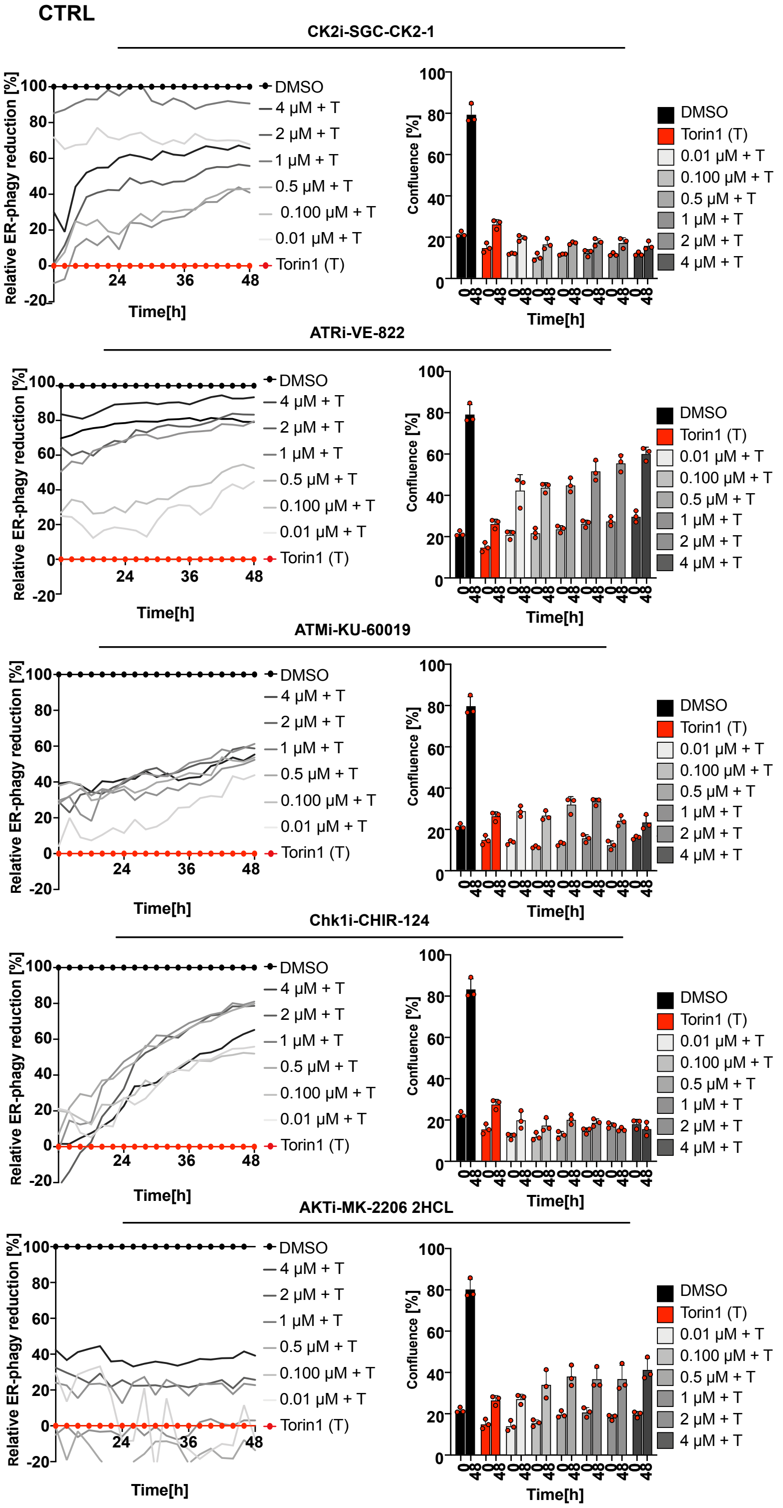


**D**

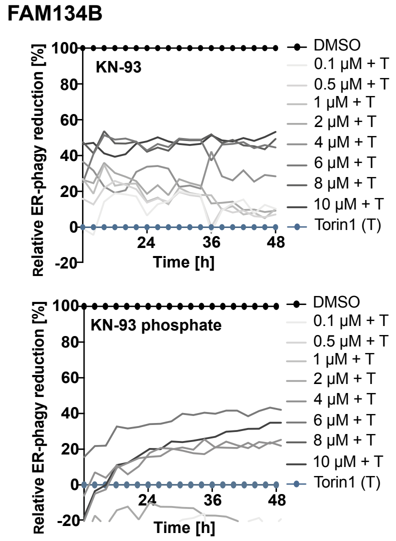
