## Supplementary material for "The function of ER-phagy receptors is regulated through phosphorylation-dependent ubiquitination pathways": Supl Fig3

### Supplementary Figure 3

**A**

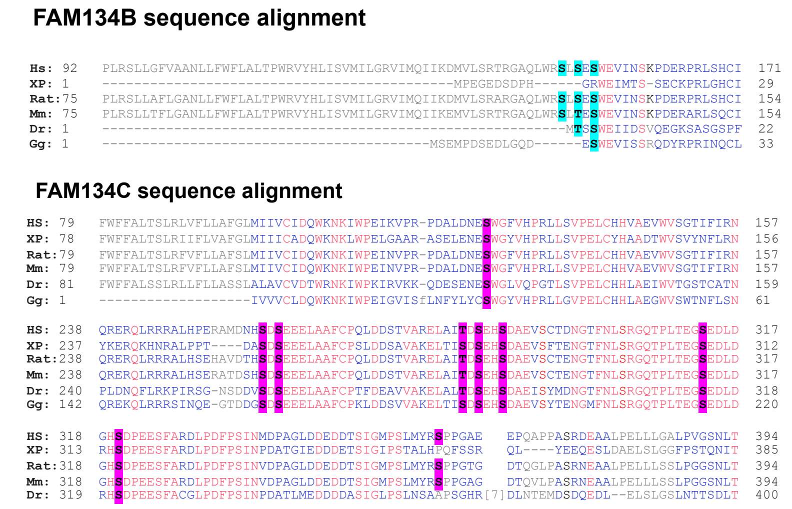


**B**

| Protein | Serine # | Predicted kinase (PHOSIDA) |
|---|---|---|
| FAM134B | S149 | **CK2** (S/T-X-X-E) GSK3 (S-X-X-X-S)) |
| | S151 | CAMK2 (R-X-X-**S/T**) **CK2** (S-X-X-**S/T**) |
| | S153 | CK1 (S/T-X-X-X-**S**) NEK6 (L-X-X-**S/T**) |
| FAM134C | S126 | **CK2** (**S/T**-X-X-E) CAMK2 (R-X-X-**S/T**, R-X-X-**S/T**-V) |
| | S258 | **CK2** (**S/T**-X-X-E) |
| | S260 | **CK2** (**S/T**-X-X-E) |
| | T283/S285/288 | **CK2** (**S/T**-X-X-E) |
| | S313 | N/A |
| | S320 | **CK2** (**S/T**-X-X-E) |
| | S360 | N/A |

**C**

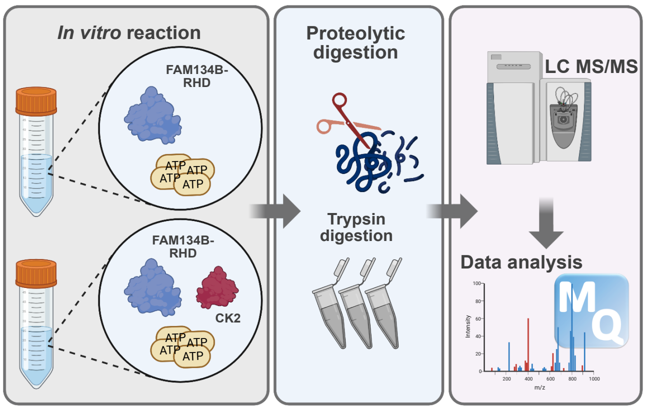


**D**

In Vitro Phosphorylation MS Results

| Positions within FAM134B wt | Protein | Phospho (STY) Probabilities |
|---|---|---|
| 153 | sp\|Q9H6L5\|His__RHD_RETR1_HUMAN | S(0.003)LS(0.028)ES(0.969)WEVINSKPDERPR |

In Vitro Phosphorylation MS Results

| Positions within FAM134B wt | Score diff | PEP | Score | Delta score | Number of Phospho (STY) |
|---|---|---|---|---|---|
| 153 | 15.3882 | 8.88E-61 | 106.36 | 102.75 | 1;2 |

In Vitro Phosphorylation MS Results

| Positions within FAM134B wt | Peptides Ck2$^-$ 01 | Peptides Ck2$^-$ 02 | Peptides Ck2$^+$ 01 | Peptides Ck2$^+$ 02 |
|---|---|---|---|---|
| 153 | | | 1 | 1 |

**E**

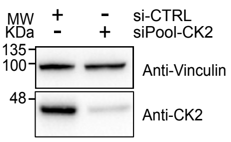


**F**

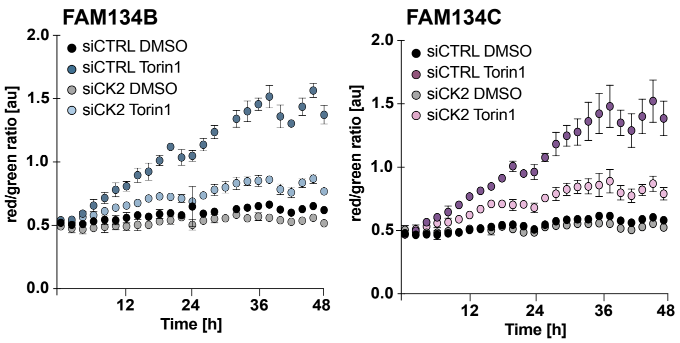
